# metaGWASmanager: A toolbox for an automated workflow from phenotypes to meta-analysis in GWAS consortia

**DOI:** 10.1101/2024.03.12.584556

**Authors:** Zulema Rodriguez-Hernandez, Mathias Gorski, Maria Tellez Plaza, Pascal Schlosser, Matthias Wuttke

## Abstract

**Summary:** This paper introduces the metaGWASmanager, which streamlines genome-wide association studies within large-scale meta-analysis consortia. It is a toolbox for both the central consortium analysis group and participating studies to generate homogeneous phenotypes, minimize variance from inconsistent methodologies, ensure high-quality association results, and implement time-efficient quality control workflows. The toolbox features a plug-in-based approach for customization.

**Results:** The metaGWASmanager toolbox has been successfully deployed in both the CKDGen and MetalGWAS Initiative consortia across hundreds of participating studies, demonstrating its effectiveness in improving the efficiency of GWAS analysis by automating routine tasks and ensuring the value and reliability of association results, thus, ultimately promoting scientific discovery.

**Availability:** GitHub: https://github.com/genepi-freiburg/metaGWASmanager

## 1 Introduction

Genome-Wide Association Studies (GWAS) aim to unravel genes implicated in health and disease and advance our understanding of physiology and pathology (Tam et al. 2019). In GWAS, obtaining meaningful results involves increasing sample sizes through meta-analysis, as exemplified by the saturation of height GWAS signals with multi-million individuals (Yengo et al. 2022). Partnership within consortia, as seen in the collaborative efforts of CKDGen over a decade (Köttgen and Pattaro 2020), is critical to achieving these sample sizes. Consortia typically establish a centralized analysis group, and study-specific analysts follow a predefined analysis plan. To ensure reliable results, maintaining high-quality, standardized phenotypic data and genotypes across studies and quality control on the contributing study summary statistics are imperative.

Therefore, we aim to offer a versatile framework that can be universally applied to accomplish complex tasks with different tools for phenotype generation and statistical analyses. This intends to minimize between-studies heterogeneity originating from the use of inconsistent methodologies, disparate data preparation, or divergent tools for statistical tests, and to save time by automating routine tasks both for the central Consortium Analysts (CA) group and the Study Analysts (SA).

In this paper, we introduce a comprehensive toolbox leveraging existing software packages and streamlining the entire consortia workflow. This includes harmonization and quality control of phenotypes, GWAS, quality control reports of GWAS (in the form of test statistics and plots) and meta-analysis, providing an integrated solution for study sites and central analysis teams (**Figure 1**).

**Figure 1.**
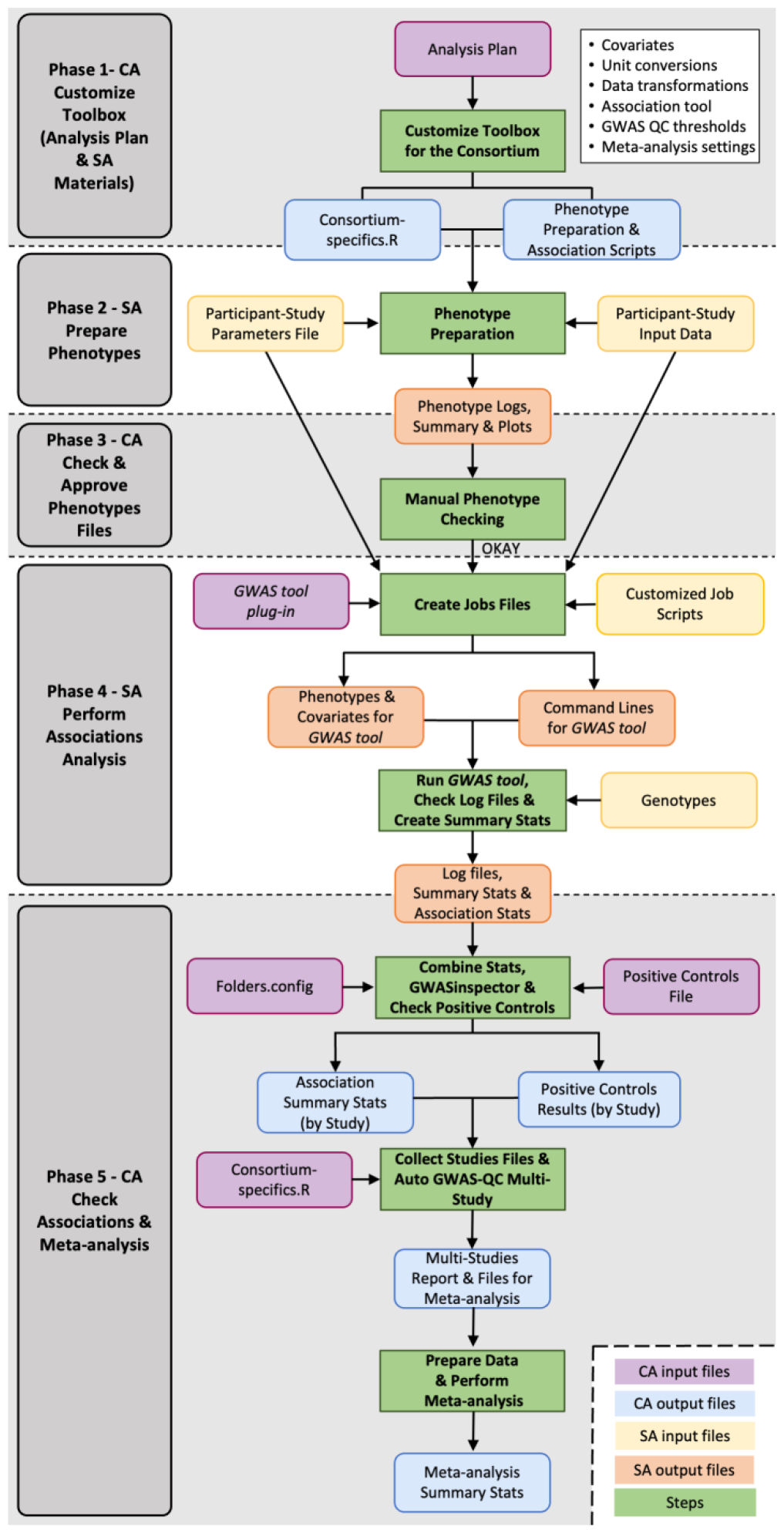
Workflow. Illustration of the metaGWASmanager pipeline, outlining phenotype generation and quality assurance, followed by the GWAS, GWAS-QC and meta-analysis steps, along with the required inputs and resulting outputs. Shaded and white sections indicate tasks to be carried out by CAs and SAs, respectively.

## 2 Methods and Workflow

For the presented tools, we build on R, Bash and Python as a foundation universally available across study sites. While CAs benefit from some familiarity with these languages to customize the toolbox to the consortium, SAs do not need specific coding skills. To fulfill the unique needs of each consortium, we adopted a plug-in concept, allowing for the flexible customization of the analysis pipeline with respect to phenotype preparation and used association software tools.

Our toolbox addresses two groups of users. First, the CA group is responsible for the analysis plan, pre- and post-GWAS quality control design, meta-analysis, and other downstream analyses. CAs do not have access to individual-level participant data. Second, the SA perform the phenotype preparation and association analyses at the study level and upload their results to the consortium server.

The workflow comprises five phases and is presented in **Figure 1**. In the first phase, starting with an analysis plan, the CAs customize the metaGWASmanager scripts according to the specific requirements of the consortium. The CAs can easily adopt the *consortium-specifics*.*R* plug-in file to meet diverse requirements aligned with the accompanying analysis plan. This enables customization of the entire analysis process, ranging from SA phenotype preparation over the actual GWAS to the centralized summary statistics quality control and meta-analysis by the CA group. MetaGWASmanager currently implements plink 2.0 (Chang et al. 2015) and regenie (Mbatchou et al. 2021) association softwares and can easily be customized for other tools. The analysis plan and the customized toolbox are distributed to the participating SAs.

In the second phase, the SAs prepare a study-specific phenotype file, using a programming language of their choice, and run the toolbox to check and summarize it. The script conducts a thorough examination of the input file, issuing warnings and detecting errors to ensure high-quality phenotypic data and avoid common pitfalls like wrong unit conversions or variations in assay methods. Furthermore, the pipeline performs necessary trait transformations, stratification, and calculations, standardizing the phenotypic data for consistency. SAs then upload the generated phenotype summary statistics (not containing individual-level data) and diagnostic plots (Supplementary Data), including descriptive graphs of trait-specific variables to the CAs.

In the third phase, CAs inspect phenotype distributions, detect outliers, and validate across studies. (**Figure 1**). After phenotype approval, in phase 4, studies proceed with the preparation of their genotypic data in a compatible format, with the help of provided scripts, and execute the GWAS. The toolbox provides job scripts and includes GWAS tool command line options to ensure best practice analysis. Results are packaged in a standardized way to facilitate CA group analyses. These include marker association statistics, phenotype summaries of the phenotypes actually used for association after sample exclusions, and diagnostic log files.

In phase 5, to maintain the highest possible data quality, the CAs employ code to properly format and verify GWAS statistics. GWAS files are examined using the GWASinspector R package (Ani et al. 2021). A wide variety of diagnostic reports are generated to facilitate the check for several potential issues, including discrepancies in genomic builds, allele switches and swaps, missing data, file and number formatting issues, unaccounted inflation, and incorrect transformations. The CA group communicates any identified issue back to the respective SA for resolution.

Lastly, our pipeline facilitates progress tracking and reporting by generating tables and plots that allow for efficient monitoring of the consortium’s progress, such as ongoing GWAS status, studies awaiting analysis, and current sample sizes. These tracking mechanisms are crucial for effective coordination and management of the consortium effort. Once data collection and QC is complete, the actual GWAS meta-analysis is conducted using METAL (Willer et al. 2010).

## 3 Applications in the CKDGen and Metals consortia

The presented toolbox has been developed in the context of the CKDGen consortium (Schlosser et al. 2021; Teumer et al. 2019; Tin et al. 2019; Tin et al. 2021; Wuttke et al. 2019) and has recently been successfully applied in an ongoing large-scale GWAS meta-analyses with over 130 participating studies of >2 million individuals with 18 endpoints and a total of >1,800 unique GWAS. We show exemplary phenotype and cleaning diagnostic plots and reports along with comprehensive documentation in the **Supplementary data**.

To prove the transferability of the workflow, we implemented the same tools with the recently established MetalGWAS Initiative studying >40 phenotypes in ∼10 studies (https://biodama.isciii.es/metal-gwas/).

Furthermore, we provide sample scripts to run the complete process using simulated data.

## 4 Conclusion

The metaGWASmanager aids to identify an increased number of associated variants by reducing between-study heterogeneity from inconsistent methodologies in phenotype preparation and divergent tools in association analyses. By establishing output file consistency, it supports the meta-analyses processes. Its technical foundation in R, Bash and Python, along with the flexible plug-in concept and comprehensive documentation, facilitates quick implementations in a wide range of settings. Moreover, it significantly streamlines analytical workflows by automating routine tasks, ultimately saving valuable time of the GWAS consortium team and making the scientific output more robust, thus facilitating biological discovery.

## Acknowledgements

We thank the members of the CKDGen core analyst group for their extended feedback on the different versions of the workflow over the past years and the GCKD study for the use of aggregate data to illustrate the application.

## References

Ani, A., et al. (2021), ‘GWASinspector: comprehensive quality control of genome-wide association study results’, Bioinformatics, 37 (1), 129–30.

Chang, CC., et al (2015), ‘Second-generation PLINK: rising to the challenge of larger and richer datasets’, GigaScience, 4(1), s13742–015

Köttgen, A. and Pattaro, C. (2020), ‘The CKDGen Consortium: ten years of insights into the genetic basis of kidney function’, Kidney Int, 97 (2), 236–42.

Mbatchou, J., et al. (2021), ‘Computationally efficient whole-genome regression for quantitative and binary traits’, Nat Genet, 53 (7), 1097–103.

Schlosser, P., et al. (2021), ‘Meta-analyses identify DNA methylation associated with kidney function and damage’, Nat Commun, 12 (1), 7174.

Tam, V., et al. (2019), ‘Benefits and limitations of genome-wide association studies’, Nat Rev Genet, 20 (8), 467–84.

Teumer, A., et al. (2019), ‘Genome-wide association meta-analyses and fine-mapping elucidate pathways influencing albuminuria’, Nat Commun, 10 (1), 4130.

Tin, A., et al. (2021), ‘Epigenome-wide association study of serum urate reveals insights into urate co-regulation and the SLC2A9 locus’, Nat Commun, 12 (1), 7173.

Tin, A., et al. (2019), ‘Target genes, variants, tissues and transcriptional pathways influencing human serum urate levels’, Nature Genetics, 51 (10), 1459–74.

Willer, C. J., Li, Y., and Abecasis, G. R. (2010), ‘METAL: fast and efficient meta-analysis of genomewide association scans’, Bioinformatics, 26 (17), 2190–1.

Wuttke, M., et al. (2019), ‘A catalog of genetic loci associated with kidney function from analyses of a million individuals’, Nat Genet, 51 (6), 957–72.

Yengo, L., et al. (2022), ‘A saturated map of common genetic variants associated with human height’, Nature, 610 (7933), 704–12.

