## Supplementary Information for "metaGWASmanager: A toolbox for an automated workflow from phenotypes to meta-analysis in GWAS consortia"

### Overview

This vignette introduces the metaGWASmanager, a comprehensive toolbox leveraging existing software packages and, streamlining the entire GWAS-consortium workflow. This encompasses the phenotype generation, quality control of phenotypes, GWAS, GWAS-QC and meta-analysis providing an integrated solution for both the participating Study Analysts (SA) and Consortium Analysts (CA). For a description, please see the corresponding manuscript: Zulema Rodriguez-Hernandez, Mathias Gorski, Maria Tellez Plaza, Pascal Schlosser* and Matthias Wuttke* (2023). "metaGWASmanager: A toolbox for an automated workflow from phenotypes to meta-analysis in GWAS consortia". under review.

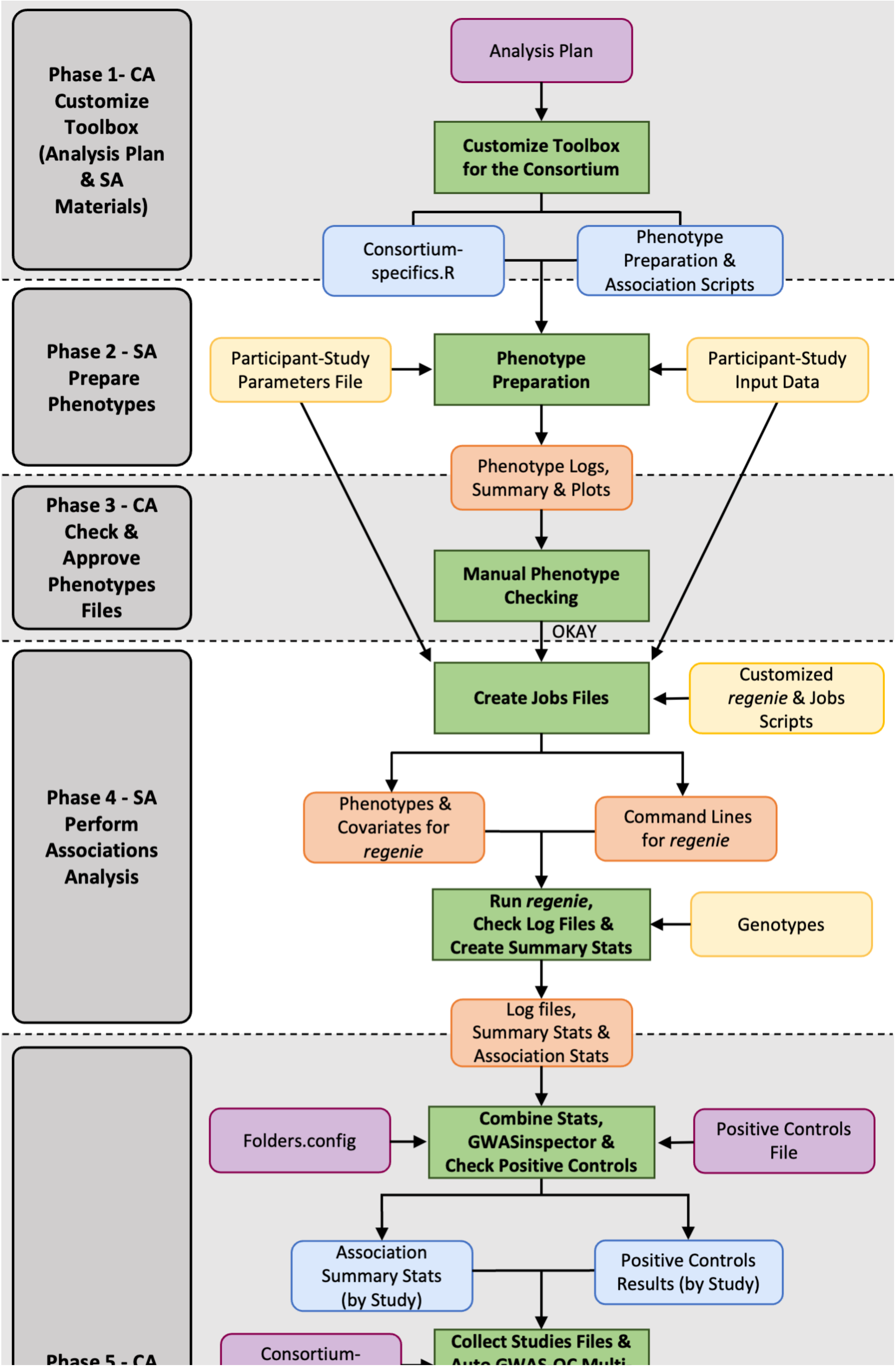
Phase 1- CA Customize Toolbox (Analysis Plan & SA Materials)
Analysis Plan
Customize Toolbox for the Consortium
Consortium-specifics.R
Phenotype Preparation & Association Scripts
Phase 2 - SA Prepare Phenotypes
Participant-Study Parameters File
Phenotype Preparation
Participant-Study Input Data
Phenotype Logs, Summary & Plots
Phase 3 - CA Check & Approve Phenotypes Files
Manual Phenotype Checking
OKAY
Phase 4 - SA Perform Associations Analysis
Create Jobs Files
Customized *regenie* & Jobs Scripts
Phenotypes & Covariates for *regenie*
Command Lines for *regenie*
Run *regenie*, Check Log Files & Create Summary Stats
Genotypes
Log files, Summary Stats & Association Stats
Folders.config
Combine Stats, GWASinspector & Check Positive Controls
Positive Controls File
Association Summary Stats (by Study)
Positive Controls Results (by Study)
Collect Studies Files &
Consortium-
Phase 5 - CA

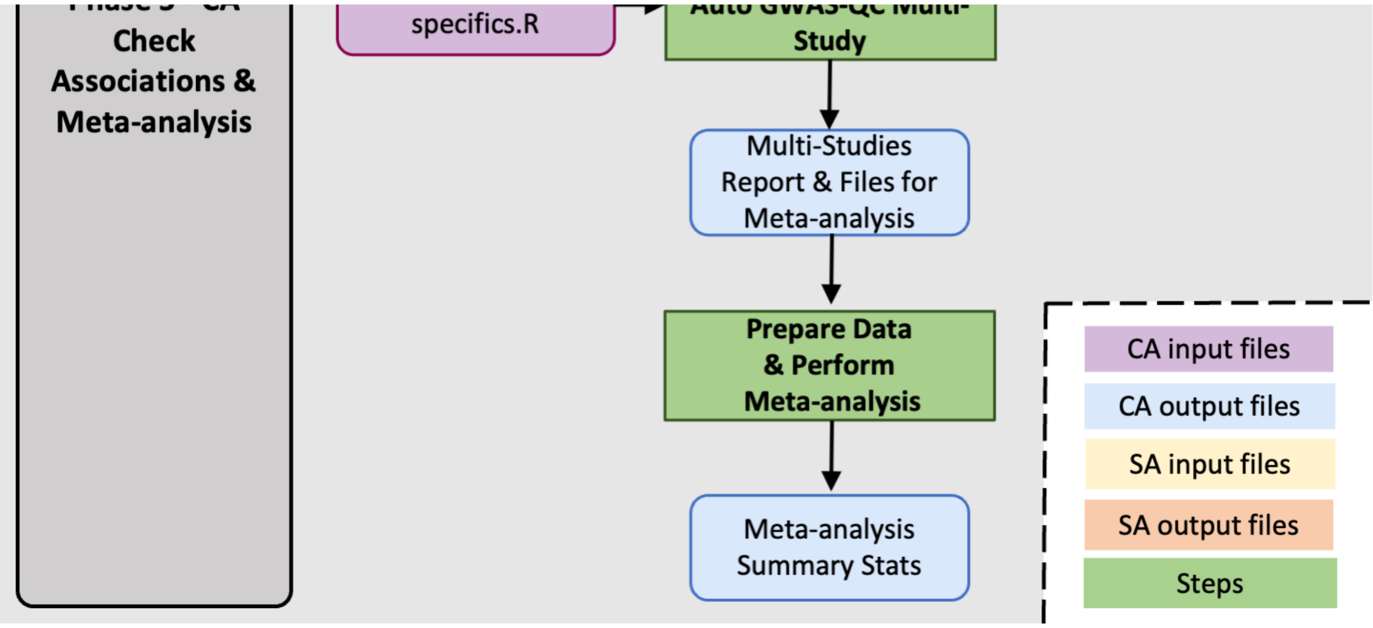


*. Illustration of the metaGWASmanager pipeline, outlining phenotype generation and quality assurance, followed by the GWAS, GWAS-QC and meta-analysis steps, along with the required inputs and resulting outputs. Shaded and white sections indicate tasks to be carried out by CA and SA, respectively.*

### Installation

To set up the toolbox please fork the github repository: https://github.com/genepi-freiburg/gwas-consortium/tree/main

### Required files, programs and server specifications

Aside from the documentation and scripts supplied by metaGWASmanager, it is necessary to ensure that the following programs and files are properly installed and downloaded on your system.

#### For SAs

- regenie
- PLINK or PLINK 2.0.
- VCFtools and BCFtools
- R software
- Analysis plan, scripts and examples files for conducting phases 2 and 4 will be provided by the CAs.

#### For CAs

Same as SAs, but also:

- GWASinspector R package
    - SQLite reference data available here
- Perl
- HTSlib

- FlexiBLAS
- Python

#### Server structure

We outline the recommended structure for the consortium server to ensure effective and efficient data management:

##### Scripts

They should be separate from data, in a different folder.

```
"/storage/consortium_name/scripts/"
```

This folder will contain the scripts needed by CAs, all provided by metaGWASmanager. It is advisable to directly download the different folders-containing scripts (*pheno_generation*, *pheno_summary*, *gwas_qc* and *metaanalysis*) from metaGWASmanager GitHub to the *scripts* folder previously created on your server:

- **pheno_generation** folder: it contains scripts needed by SAs to prepare phenotype data, and perform GWAS. To be used by SAs.
- **pheno_summary** folder: it contains scripts to summarize phenotype submissions across studies. For CAs' use.
- **gwas_qc** folder: it contains scripts to summarize genotype submissions across studies. For CAs' use.
- **metaanalysis** folder: it contains scripts to prepare data and perform meta-analysis. For CAs' use.

Please, do not modify the scripts names.

##### Pheno Upload

The phenotypes results, output of **Phase 2 - SA: Prepare Phenotypes (SAs)**, submitted by the SAs will be stored in the *uploads/pheno* directory. This directory will contain a folder for each study that has submitted the phenotype summary statistics.

```
"/storage/consortium_name/uploads/pheno/study_name"
```

Besides the study-specific folders, it should also contain two additional directories:

- *00_ARCHIVE*. If a specific-study re-upload the phenotype summary statistics, perhaps due to the detection of an error, CAs will archive the previous version in this directory.
- *00_SUMMARY*. The outcomes from performing cross-study phenotype summaries in the **Phase 3 - CA: Check & Approve Phenotypes Files** will be stored in this location.

##### GWAS Upload

The GWAS results (output of **Phase 4 - SA: Perform Associations Analysis**) submitted by the SAs will be stored in the *uploads/assoc* directory. This directory will contain a folder for each study that has submitted genetic summary statistics.

```
"/storage/consortium_name/uploads/assoc/study_name"
```

Similar to the *uploads/pheno* directory, it is recommended to create a *00_ARCHIVE* sub-directory.

It is advised to maintain consistency in the naming of studies withing both the *uploads/pheno* and *uploads/assoc* folders to prevent potential problems when cross-referencing files between the two folders.

###### Cleaning

Contains intermediate files, logs and summary files resulting from **Phase 5 - CA: Check Associations & Meta-analysis**.

```
"/storage/consortium_name/cleaning/study_name"
```

In addition to the folders designated for each study, it should also encompass two extra directories:

- *00_ARCHIVE*. To save older results.
- *00_SUMMARY*. It will contain the output of cross-study GWAS verification process (**Phase 5 - CA: Check Associations & Meta-analysis**) including plots, overall statistical summaries, positive controls validation result, among others.

Note that, the sub-folder "*data*" withing each study folder contains the *regenie* results combined by chromosomes.

```
"/storage/consortium_name/cleaning/study_name/data"
```

###### Meta-analysis

Finally, the metaanalysis folder will contain as many subfolders as target traits that CAs will create (e.g, uacr_int trait).

```
"/storage/consortium_name/metaanalysis/uacr_int"
```

### Step by step guide

#### Phase 1 - CA: Customize Toolbox (Analysis Plan & SA Materials)

The initial stage involves generating Consortium-specific files (*consortium-specifics.R* and *parameters.txt* plug-in) and formulating the Analysis Plan, in which CAs will define, based on the provided metaGWASmanager examples, the required settings for conducting the pool-cohort GWAS.

A) ***consortium-specifics.R***. The *consortium-specifics.R* plug-in file should be edited according to the CA specifications (located in the *pheno_generation* folder on the metaGWASmanager GitHub repository). This file contains several functions such as unit conversion, traits transformations, verification of the input and parameteres files, setting quality control parameters, determination of covariates, stratification etc. These functions will be applied in subsequent stages of the workflow process. The *consortium-specifics.R* file will be provided, along the Analysis Plan and required scripts, to each SAs.

B) **Parameters file**. CA will customize the *parameters.txt* file to match the particular traits to study. It must included important points, which must be filled out by the SAs, such as the name of the input file and the participant-study, SAs contact information, the study's ancestral background, the analysis date, the units of the variables to be studied, along with laboratory particular settings (e.g., limits of detection), the imputation reference panel, the number of genetic principal components, and some more optional fields. A parameters file example is provided within the *pheno_generation* folder on the metaGWASmanager GitHub repository.

C) **Analysis Plan**. The Analysis Plan PDF document should provided comprehensive information for SAs on to proceed with the different analyses, including **Phase 2 - SA: Prepare Phenotypes** and **Phase 4 - SA: Perform Associations Analysis**. Certainly, you are welcome to use the information described in this documentation related to these phases. It is also essential to include CAs contact details in case any questions arise.

#### Phase 2 - SA: Prepare Phenotypes

In a first step, CA will ask for preparation the phenotypic data for all participating studies. CA will supply an Analysis Plan and the *pheno_generation* folder.

Input files for each participating study, include an input data (***input.txt***) and a parameter file (***parameters.txt***) that SAs will create and complete, respectively, according the CA guide standards.

After that, SAs will execute the ***01-phenotypes-generation.sh*** script, which automatically generates the descriptive statistics of phenotypes, on a Linux server as follows:

```
bash 01-phenotypes-generation.sh parameters.txt
```

Note that if SAs prepare their data in a Windows or Mac system, to avoid problems with line breaks when running the script in Linux/Unix, SAs should use the following commands respectively to convert the files prior to running the script:

```
dos2unix input.txt
```

```
mac2unix input.txt
```

During this initial phase, a comprehensive examination of the input and parameters files will be carried out. This involves issuing warnings and identifying errors to guarantee high-quality phenotypic data and prevent common pitfalls such as unit conversion discrepancies or variations in assay methods. Additionally, the pipeline executes essential trait transformations and calculations, ensuring uniformity in the phenotypic data.

The files generated by phenotype preparation scripts include the phenotype summary statistics (***STUDYNAME_ids_summary.txt***) and plots files. SAs must carefully examine the *.log* files for errors or warnings as well as whether the generated descriptive statistics and plots are reasonable in the pdf file. If there are problems, SAs will try to adjust the parameters in the ***parameters.txt*** or ***input.txt*** file according to the different error messages.

In case everything seems fine, SAs will send all the files from the generated *return_pheno* folder via email to CAs. The CAs will then check phenotype descriptive data. It is important to note that the files uploaded by SAs contain only summarized statistical data and never the original information from study participants.

You can find an example of a phenotype generation report here

#### Phase 3 - CA: Check & Approve Phenotypes Files

The submited files within the **return_pheno** folder will be stored in the *upload/pheno* directory by CAs, within the folder corresponding to the specific participating study.

```
"/storage/consortium_name/uploads/pheno/study_name"
```

Subsequently, CA will manually inspect these files and plots in order to identify potential issues and inconsistencies. Furthermore, CA will run the scripts located in the **pheno_summary** directory of the metaGWASmanager toolkit (***01_collect_summary.sh*** and ***02_plot_summaries.sh***).

Both scripts summarize phenotype summary-statistics submissions (***STUDYNAME_ids_summary.txt***) across studies and also facilitate their representation for improved inter-study comparison and to detect potential outliers. Please run both scripts using the *bash* command line from:

```
"/storage/consortium_name/scripts/pheno_summary"
```

Note that the generated outputs of the inter-study comparison analyses will be automatically saved in the *00_SUMMARY* folder located at:

```
"/storage/consortium_name/uploads/pheno/00_SUMMARY"
```

After the inspection of the submitted files, CA will provide feedback to SAs. In the case CA detects an issue, it will be clearly outlined and assistance will be offered to the SAs in oder to resolve it. The SAs will then revisit **Phase 2 - SA: Prepare Phenotypes** and re-submit the corresponding outputs files. Contrary, if no problems are identified, SAs are authorized to proceed with **Phase 4 - SA: Perform Associations Analysis**.

#### Phase 4 - SA: Perform Associations Analysis

To initiate this step, SAs should have received the approval from CA. Additionally, they must have prepared the measured genotype (chip data) and imputed data for *regenie* steps 1 and 2 following the instructions provided in the Analysis Plan. More information about *regenie* steps here.

Please note that *regenie*-step1 needs the measured genotype data in PLINK format. We recommend to extract the high quality variants following the *regenie* best practices:

```
plink2 --geno 0.1 --hwe 1e-15 --mac 100 --maf 0.01 --mind 0.1 --indep-pairwise 1000
100 0.9
```

For that, SAs can use the provided ***plink_qc.sh*** script.

In addition, *regenie*-step2 requires the imputed genetic data (chunked by chromosome) in *.bgen* format v1.2 with 8 bits encoding. SAs can easily convert the imputed data to *.bgen* format using the provided ***vcf_to_bgen.sh*** script, inside the *pheno_generation* folder or by running the following commands on Linux:

```
plink2 --vcf $infile dosage=DS --make-pgen erase-phase --out $pgen_unphased
```

```
plink2 --pfile $pgen_unphased --export bgen-1.2 bits=8 --out $out_bgen
```

Once the genetic data is prepared, the SAs can systematically initiate the necessary script executions.

A) **Create job files**.

First, SAs should adjust the following fields of ***make-regenie-step1-job-scripts.sh*** and ***make-regenie-step2-job-scripts.sh*** scripts according to their *regenie*-installation, data pathway and prefix of their genotyped and imputed files.

For ***make-regenie-step1-job-scripts.sh***:

```
. REGENIE=<path to your regenie software>
. PLINK_DATA_PREFIX=<path and prefix to your PLINK *.bed file>
. PLINK_SNP_QC=<path to your *.snplist file containing a list of high quality variants used for
regenie-step1>
. PLINK_INDIV_QC=<path to your *.id file containing a list of high-quality individuals used for
regenie-step1>
. --threads <Number of threads that should be utilized by each individual regenie-step1 job>
```

For ***make-regenie-step2-job-scripts.sh***:

```
. REGENIE=<path to your regenie software>
. BGEN_SAMPLE_PREFIX=<path to your *.bgen and *.sample files> (as one sample file exists per
chromosome; please adjust the prefix file name using ${CHR} as placeholder for the chromosome).
. --threads <Number of threads that should be utilized by each individual regenie-step-2 job>
```

Note that these scripts are set up to utilize a Slurm job scheduler.

Upon making the mentioned adjustments, SAs can executed the ***02-make-regenie-jobs.sh*** script.

```
bash 02-consortium-make-regenie-jobs.sh parameters.txt
```

The process will generate the phenotypes and covariates details along with the necessary command lines to run *regenie*-steps. As a result, in the *jobs* folder, for *regenie*-step1, as many jobs as phenotype will be created, and one job for each phenotype and chromosome for *regenie*-step2.

B) **Run GWAS using *regenie***

SAs then will execute ***03-submit-all-jobs.sh*** to submit all GWAS according to the jobs generated in the preceding step for both *regenie*-steps 1 and 2.

```
bash 03-submit-all-jobs.sh
```

Our recommendation is to first run a pilot analysis (e.g., one phenotype for *regenie*-step1, one chromosome for *regenie*-step2) before submitting all jobs.

C) **Create Summary Statistics** and **Collect & Upload results**.

The SAs will then run the ***04-postprocess-results.sh*** script, which investigate the different log files generated during the *regenie* process to ensure the successful execution of each GWAS. It also produces tailored summary tables for each association analysis.

```
bash 04-postprocess-results.sh
```

If an issue occurs, SAs should investigate the cause. After resolving it, SAs can re-execute individual phenotypes and/or steps by using the appropriate files from the *jobs* directory. In case SAs needed to change any parameter o paths, they would have to re-run the ***02-consortium-make-regenie-jobs.sh*** again.

Finally, the results will be compiled into a compressed folder using ***05-collect-files-for-upload.sh***

```
bash 05-collect-files-for-upload.sh study_name
```

It will be created a a gzipped tar-ball folder containing outputs from *return_pheno* folder including log files, summary tables and plots, and the *regenie*-step2 outputs. The compressed folder will then be submitted to the CA server for further validation.

We also recommend that CAs, at this point, somehow collect information about the specific authors (name of authors and affiliations, conflict of interest, etc) and studies (genotyping information, study acknowledgements,relevant study information). For example, CAs may use a Google Sheets and/or Google Forms.

#### Phase 5 - CA: Check Associations & Meta-analysis

The compressed folder submitted by the SAs, will be stored in the *upload/assoc* directory, within the folder corresponding to the specific participating study.

```
"/storage/consortium_name/uploads/assoc/study_name"
```

CA will then proceed to unzip the folder and run scripts located in the *gwas_qc* folder of the metaGWASmanager toolbox. Please, note that scripts located in *gwas_qc* will be run directly from this folder.

```
"/storage/consortium_name/scripts/gwas_qc"
```

###### ***folders.config.sh*** file

It includes details about essential pathways (the script folder, cleaning directory, upload data, etc) and specific variables (phenotypes, ancestry and strata vectors, among others) needed for executing downstream scripts. CAs first should customize these fields and may also incorporate important settings to align with particular server requirements. The ***folders.config.sh*** file will be sourced throughout subsequent scripts, so no modification to other scripts are necessary.

After making the necessary settings to the *folders.config.sh* file, CA are ready to carry out the subsequent steps.

A) ***01_combine_chromosomes.sh***.

It will check if any file are missing from the *output_regenie_step2* folder (one file for each phenotype and chromosome) ensuring the presence of the required columns using *find-column-index.pl* script. Once the validation process is successfully finalized, it merge all chromosomes into a single file for each trait. Subsequently, each file is then compressed and a corresponding tabix file is generated.

Command line to run:

```
bash 01_combine_chromosomes.sh study_name
```

B) ***02_gwasinspector.sh***.

CAs will then run GWASinspector:

```
bash 02_gwasinspector.sh study_name
```

GWASinspector performs an extensive quality control process on GWAS results ensuring their accuracy, proper format, and consistency across all studies using robust criteria. As a result, in the *cleaning/study_name/qc_output* folder, clean and harmonized files will be created along with a comprehensive report and plots (including Manhattan and QQ plots, histograms, etc) as shown below:

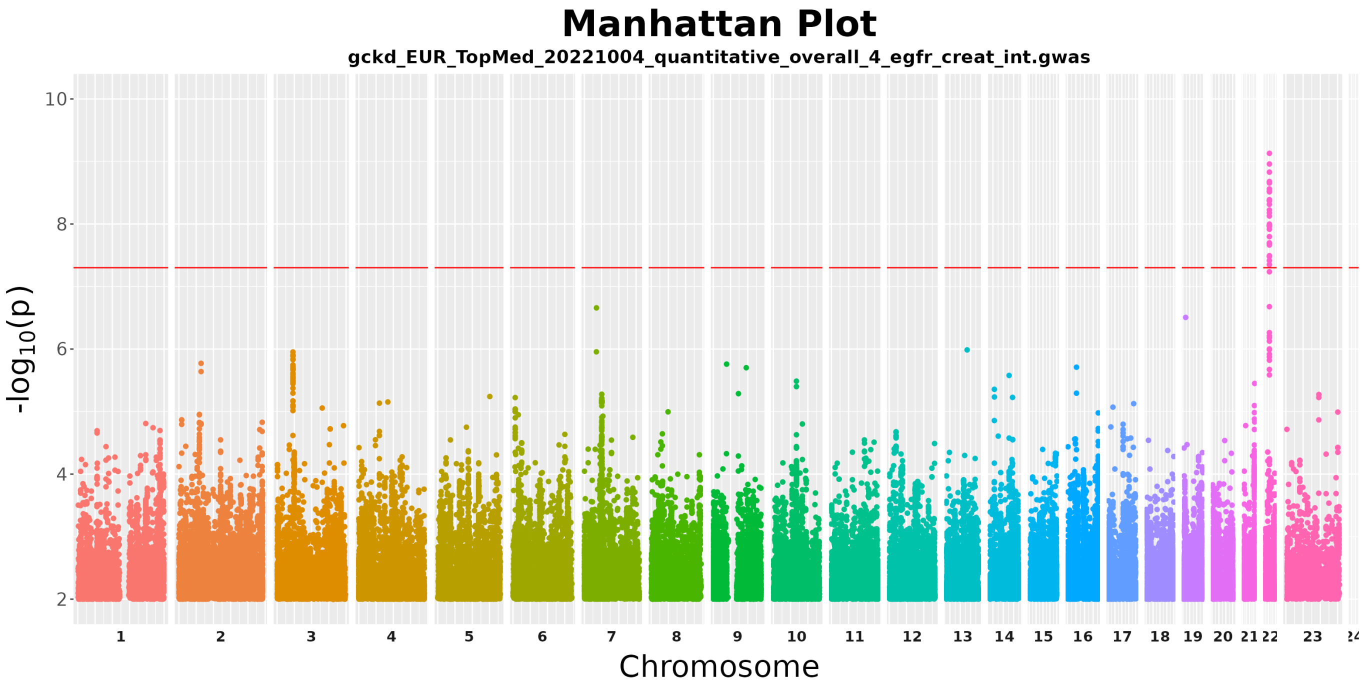


*Manhattan plot created by GWASinpector workflow of overall eGFR creatinine in the GCKD cohort, a participating study in the CKDGenR5 consortium.*

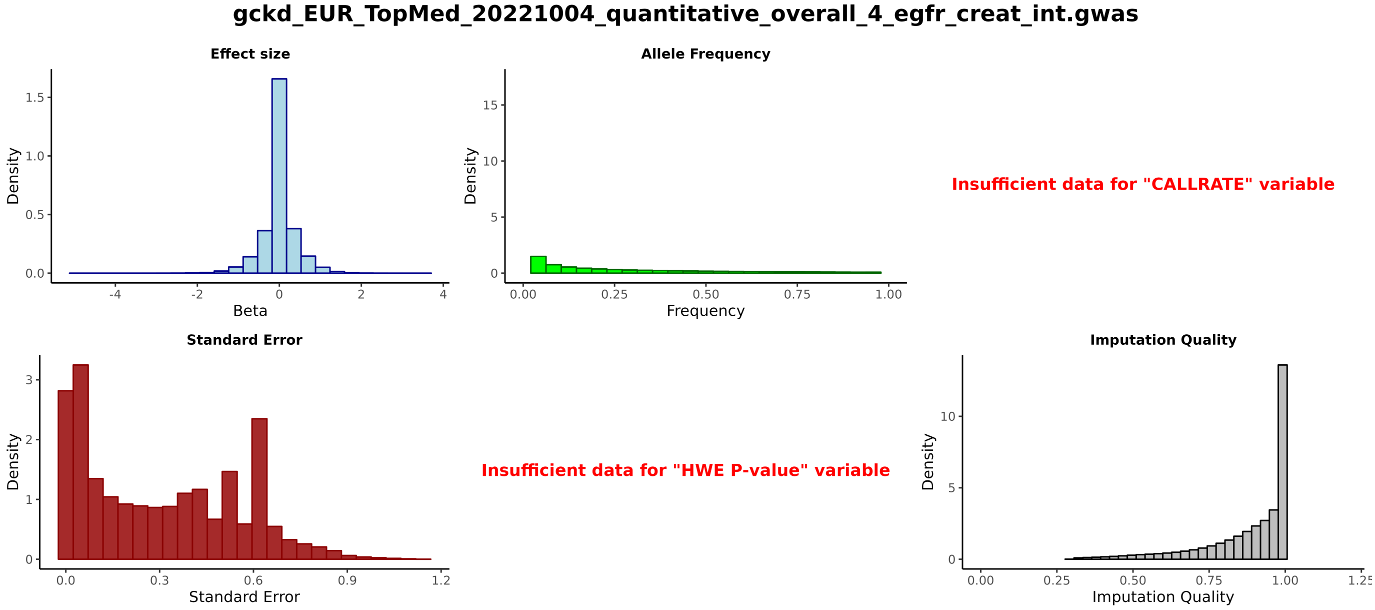


*Quality controls plot created by GWASinpector workflow of overall eGFR creatinine in the GCKD cohort, a participating study in the CKDGenR5 consortium.*

More information about GWASinspector R package here.

C) ***03_check_positive_controls.sh***

CAs should create a GWAS positive control *.txt* file containing the hits (specific to each target trait and ancestry) widely validated in the literature, whenever possible.

CAs can also find an example in the *gwas_qc/Positive_Controls* folder (please, replace the provided example by your own customized ***positive-controls.txt*** file):

| trait | ethnicity | SNP | gene | chr | pos_b37 | pos_b38 | ref | alt | MA |
|---|---|---|---|---|---|---|---|---|---|
| egfr_creat_int | EUR | rs13329952 | UMOD | 16 | 20366507 | 20355185 | T | C | 0.1 |
| egfr_creat_male | EUR | rs13329952 | UMOD | 16 | 20366507 | 20355185 | T | C | 0.1 |
| egfr_creat_female | EUR | rs13329952 | UMOD | 16 | 20366507 | 20355185 | T | C | 0.1 |
| egfr_creat_int | EAS | rs10277115 | UNCX | 7 | 1285195 | 1245559 | A | T | 0.3 |
| egfr_creat_male | EAS | rs10277115 | UNCX | 7 | 1285195 | 1245559 | A | T | 0.3 |
| egfr_creat_female | EAS | rs10277115 | UNCX | 7 | 1285195 | 1245559 | A | T | 0.3 |

By running ***03_check_positive_controls.sh***, each file within the *cleaning* directory will be checked and compared with the created ***positive-controls.txt***

```
bash 03_check_positive_controls.sh study_name
```

Results will be saved in ***03-check-positive-controls.csv*** file withing the corresponding folder:

```
"/storage/consortium_name/cleaning/study_name"
```

D) ***04_collect_qc_stats.sh***

It will check all clean studies after GWASinspector quality control:

```
"/storage/consortium_name/cleaning/study_name/qc_output"
```

and generate a ***qc-stats.csv*** and ***qc-stats.xlsx*** files in the *00_SUMMARY* folder:

```
"/storage/consortium_name/cleaning/00_SUMMARY"
```

The ***qc-stats.csv*** contains information about the specific study, phenotype, ancestry, sample size, variants per chromosome, lambda and summary stats for both "All" and "High quality" variants such as P-value (PVAL), effect allele frequency (EFF_ALL_FREQ), imputation quality (IMP_QUALITY), effect size (BETA), standard error (STDERR). The file then, will contain as many rows as specific studies and phenotypes.

| STUDY | PHENO | POP | PVALUE_MED_ALL | EFF_ALL_FREQ_MED_ALL | IMP_QUALITY_MED_ |
|---|---|---|---|---|---|
| GCKD_2022-10-04 | ckd | EUR | 0.4822 | 0.001215 | 0.9469 |
| GCKD_2022-10-04 | ma | EUR | 0.4787 | 0.001851 | 0.9572 |
| GCKD_2022-10-04 | gout | EUR | 0.4743 | 0.001198 | 0.9466 |

Command line to run:

```
bash 04_collect_qc_stats.sh
```

In addition, this script generates a general (all specific studies) positive controls summary files (***positive-controls.csv*** and ***positive-controls.xlsx***) in the *00_SUMMARY* folder.

E) ***05_plot_qc_stats.sh***

Multi-studies summary plots will be then generated from ***qc-stats.csv*** file within the *00_SUMMARY/plots* folder:

```
"/storage/consortium_name/cleaning/00_SUMMARY/plots"
```

Command line to run:

```
bash 05_plot_qc_stats.sh
```

F) ***06_plot_frequencies.sh***

Running ***06_plot_frequencies.sh*** generates a correlation plot for each phenotype across all the specific studies. The objective is to verify the correlation of allele frequencies between the observed data (a phenotype of specific study) with a reference panel.

```
bash 06_plot_frequencies.sh
```

Plots will be save in study-specific folder.

```
"/storage/consortium_name/cleaning/study_name/freqs"
```

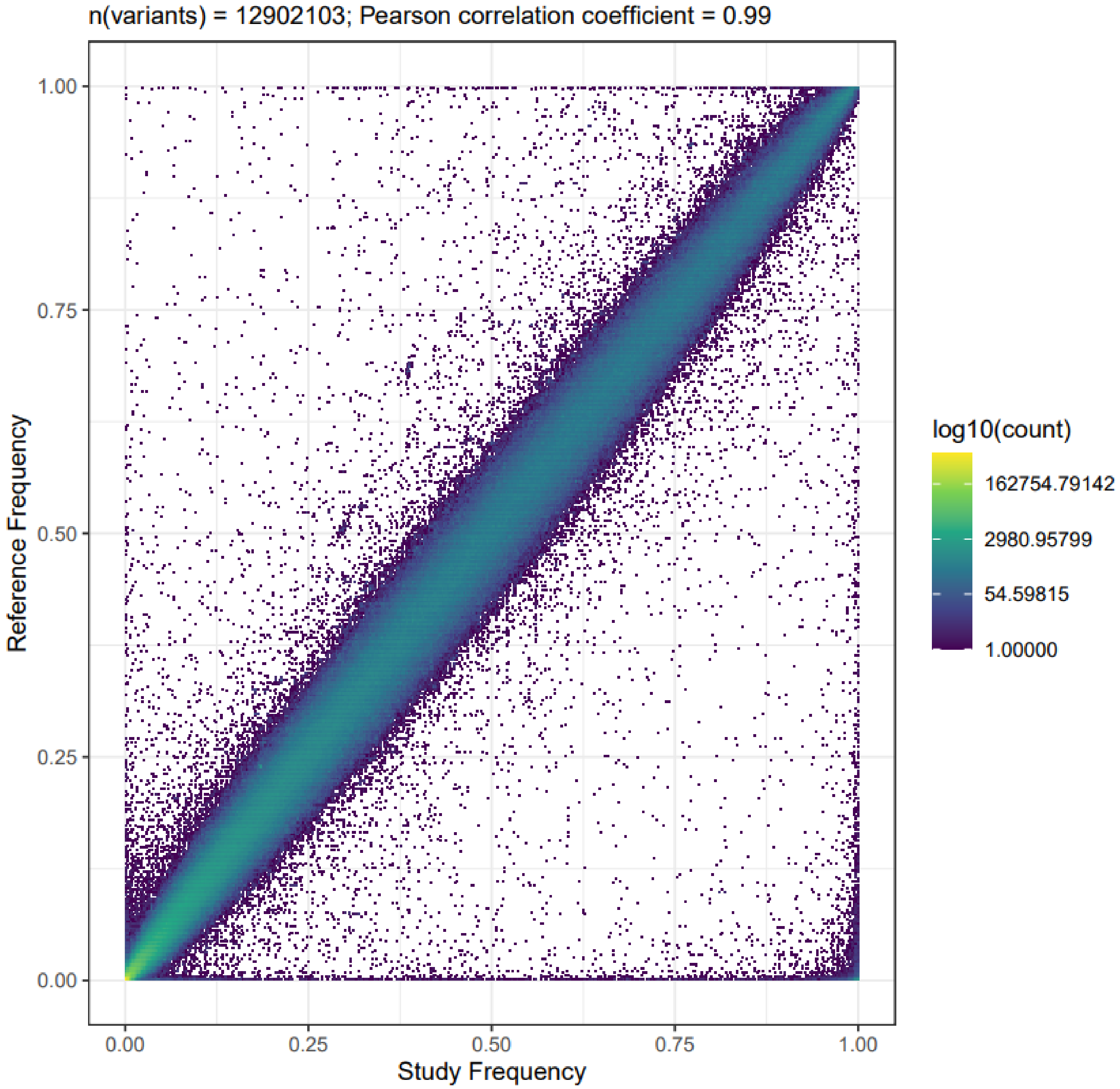


*Frequency correlation plot of overall eGFR creatinine in the GCKD cohort, a participatin study in the CKDGenR5 consortium.*

G) ***07_stateOfAffairs.sh***

Efficient monitoring of the consortium's progress, such as ongoing GWAS status, studies awaiting analysis, and current sample sizes is crucial for effective coordination and management of the consortium effort. The ***07_stateOfAffairs.sh*** print such report and a pie plot from ***qc-stats.csv*** file.

Command line to run:

```
bash 07_stateOfAffairs.sh
```

H) ***08_checkAllFileNames.sh***

For proper execution of subsequent steps (Auto GWAS quality control and meta-analysis) a final check is important to ensure consistency in files names, studies and phenotypes. The ***09_checkAllFileNames.sh*** scripts systematically validate these aspects and report if any inconsistencies arise.

```
bash 09_checkAllFileNames.sh
```

I) ***09_GWAS_QC_multistudies.sh***

By combining the summary statistics from GWAS QC (***qc-stats.csv***) and the results of positive controls analysis (***positive-controls.csv***) across all participating studies, CA will perform a final checking process. A set of relevant items will be checked, including: wrong genomic build, allele switches and swaps, missing data, file and number formatting issues, unaccounted inflation, and wrong transformations, among others.

metaGWASmanager provides a script (***10_GWAS_QC_multistudies.sh***), which automatically evaluates the list of potential problems, mentioned above. For its execution, CA will require the previously edited ***consortium-specifics.R*** plug-in, located in the *pheno_generation* folder, which supplies a tolerance table, establishing the thresholds to apply at each verification stage based on the different phenotypes and covariates to be tested.

Command line to run:

```
bash 10_GWAS_QC_multistudies.sh
```

In the end, a conclusive-summary table will be generated in the *cleaning/00_SUMMARY/AutoQC* folder, indicating whether each trait for each assessed item is categorized as "OKAY" or "NOT OKAY".

| study | pheno | pop | N | pval | lambda | impqual | beta | stderr | allel_ |
|---|---|---|---|---|---|---|---|---|---|
| GCKD_2022-10-04 | ckd | EUR | 4993 | OK | OK | OK | OK | OK | OK |
| GCKD_2022-10-04 | ma | EUR | 3998 | OK | NOT OK | OK | OK | OK | OK |
| GCKD_2022-10-04 | gout | EUR | 5034 | OK | NOT OK | OK | OK | OK | OK |
| GCKD_2022-10-04 | albumin_serum_int | EUR | 4992 | OK | OK | OK | NOT OK | OK | OK |
| GCKD_2022-10-04 | egfr_creat_int | EUR | 4993 | OK | OK | OK | OK | OK | OK |

CA should scan all entries and emphasizing those labeled as "NOT OKAY", as well as, plots produced by GWASinspector step. If an explanation for any identified issues cannot be determined, CAs must reach out to the SAs to collaboratively work towards a resolution.

After completing all GWAS quality controls checks, CAs are pepared to proceed with the meta-analysis pipeline. The necessary scripts are located in the *metaanalysis* folder provided by metaGWASmanager, and they should be run here:

```
"/storage/consortium_name/scripts/metaanalysis"
```

J) ***01-locate-input-files.sh***

First, CAs must prepare the ***input-file-list.txt*** containing all the study names to be meta-analyzed as follows:

| |
|---|
| GCKD_2022-10-04 |
| Study_name_2 |
| Study_name_3 |
| ... |
| Study_name_N |

The file should be save withing the created *metaanalysis* directory.

```
"/storage/consortium_name/metaanalysis"
```

Given a phenotype (e.g., uacr_int) the ***01-locate-input-files.sh*** script creates the ***input-files-with-path.txt*** file with the full paths to the cleaned *regenie*-step2 files for the specific phenotype across all studies intended for meta-analysis.

```
bash 01-locate-input-files.sh uacr_int
```

K) ***02-prepare-ma-input.HQ.sh*** and ***02-prepare-ma-input.LQ.sh***

They are both mostly the same scripts, but the first with stringent filter settings and second with lenient filter settings. Based on the filter configurations, input files per specific phenotype and study will be generated in the *input* folder.

```
"/storage/consortium_name/metaanalysis/uacr_int/HQ/input"
```

Command line to run:

```
bash 02-prepare-ma-input.HQ.sh uacr_int
```

L) ***03-prepare-metal-params.sh***

It creates the ***metal-params.txt*** file needed to perform meta-analysis. CAs can find the generated file at:

```
"/storage/consortium_name/metaanalysis/uacr_int/HQ/metal_output_HQ"
```

To run the script, the phenotype (e.g., uacr_int) and type of selected variants (e.g, LQ or HQ) must be provided as arguments:

```
bash 03-prepare-metal-params.sh uacr_int HQ
```

M) ***04-run-metal.sh***

Finally, run meta-analysis as follow:

```
bash 04-run-metal.sh uacr_int HQ
```

Results will be stored in a **output** folder:

```
"/storage/consortium_name/metaanalysis/uacr_int/HQ/metal_output_HQ/output"
```

Here CAs will find a **.info** file, encompassing details about the meta-analysis process, parameters used and input files. Aditionally a *.tbl* file will be available, containing the meta-analysis results.

| MarkerName | Allele1 | Allele2 | Freq1 | FreqSE | MinFreq | MaxFreq | Effect | St[illegible] |
|---|---|---|---|---|---|---|---|---|
| 3:53066069:C:G | c | g | 0.9993 | 0.0000 | 0.9993 | 0.9993 | -2.59795000 | 3.278[illegible] |
| 3:100796605:A:G | a | g | 0.9936 | 0.0003 | 0.9929 | 0.9937 | 0.10832363 | 0.101[illegible] |
| 1:174816432:C:T | t | c | 0.0051 | 0.0015 | 0.0041 | 0.0073 | -0.08297466 | 0.124[illegible] |
| 7:108043353:A:C | a | c | 0.0002 | 0.0000 | 0.0002 | 0.0002 | 0.37409000 | 0.639[illegible] |
| 2:234097095:A:G | a | g | 0.1593 | 0.0128 | 0.1519 | 0.1814 | -0.00944623 | 0.021[illegible] |
| 5:128938723:G:T | t | g | 0.2990 | 0.0011 | 0.2968 | 0.2995 | -0.00445147 | 0.017[illegible] |
| 8:2498343:C:T | t | c | 0.0002 | 0.0000 | 0.0002 | 0.0002 | 0.17083500 | 0.642[illegible] |

More information about the meta-analysis process here.
